# Partial FMRP expression is sufficient to normalize neuronal hyperactivity in Fragile X neurons

**DOI:** 10.1101/608331

**Authors:** John D. Graef, Hao Wu, Carrie Ng, Chicheng Sun, Vivian Villegas, Deena Qadir, Kimberly Jesseman, Stephen T. Warren, Rudolf Jaenisch, Angela Cacace, Owen Wallace

**Author notes:** Correspondence should be addressed to John D. Graef and Hao Wu. equal contribution (John D. Graef, Hao Wu).

## Abstract

Fragile X Syndrome (FXS) is the most common genetic form of intellectual disability caused by a CGG repeat expansion in the 5’-UTR of the Fragile X mental retardation gene *FMR1*, triggering epigenetic silencing and the subsequent absence of the protein, FMRP. Reactivation of FMR1 represents an attractive therapeutic strategy targeting the genetic root cause of FXS. However, largely missing in the FXS field is an understanding of how much *FMR1* reactivation is required to rescue FMRP-dependent mutant phenotypes. Here, we utilize FXS patient derived excitatory neurons to model FXS *in vitro* and confirm that the absence of FMRP leads to neuronal hyperactivity. We further determined the levels of FMRP and the percentage of FMRP positive cells necessary to correct this phenotype utilizing a mixed and mosaic neuronal culture system and a combination of CRISPR, antisense and expression technologies to titrate FMRP in FXS and WT neurons. Our data demonstrate that restoration of greater than 5% of overall FMRP expression levels or greater than 20% FMRP expressing neurons in a mosaic pattern is sufficient to normalize a FMRP-dependent, hyperactive phenotype in FXS iPSC-derived neurons.

**Highlights:**

- CRISPR gene editing to generate FMRP KO and CGG-deleted isogenic iPSCs
- MEA as an approach to identify FMR1 dependent phenotype in NGN2 neurons derived from FXS and FMRP KO iPSCs
- Cell mixing paradigm as mosaicism in a dish to rescue phenotype
- Minimal level of FMRP determined by FMR1 mRNA and targeted demethylation of CGG repeats to correct the hyperactive phenotype in FXS neurons
- ASO titration-validated partial expression of FMRP is sufficient to normalize increased neuronal activity

## Introduction

Fragile X Syndrome (FXS) is a neurodevelopmental disorder caused by a CGG repeat expansion in the 5’-UTR of the Fragile X mental retardation gene, *FMR1* [1]. Expansions of >200 repeats can lead to hypermethylation of the CGG repeats and CpG islands in the upstream promoter region. This hypermethylation results in heterochromatin formation and silencing of the *FMR1* transcript, thereby preventing FMRP protein production [2,3]. FMRP is an RNA-binding protein [4,5] that is highly expressed in neurons [6,7] where it plays a key role in regulating local activity-dependent synaptic translation [8]. The absence of FMRP affects both synaptic formation and maturation [9], as well as different forms of synaptic and homeostatic plasticity [10–13]. A potential manifestation of this abnormal synaptic function is increased excitability in FXS neurons [14]. For example, increased seizure susceptibility has been observed in both FXS patients [15] and *Fmr1* knockout mice [16]. Moreover, studies in *Fmr1* knockout mice showing aberrant ion channel expression and function [17–20], altered intrinsic neuronal properties [21,22] and augmented network activity [22,23] all demonstrate FMRP-dependent effects on neuronal hyperexcitability. The *Fmr1* knockout mice have been a good model for understanding the signaling pathways, pathophysiology, and behavioral phenotypes associated with FXS. However, disease-modifying therapeutics from mouse models have not translated well to the clinic [24].

FXS patient derived iPSCs represent an alternative model system to identify potential strategies for *FMR1* reactivation, rather than targeting downstream pathways. FXS iPSCs retain expanded CGG repeats, promoter hypermethylation and FMR1 silencing after reprogramming [25,26], and have been used to demonstrate that removal of the expanded CGG repeat region leads to demethylation of the FMR1 promoter and reactivation of *FMR1* [27,28]. Additionally, recent studies have shown that removal of the CGG repeat region in FXS iPSC-derived neurons not only fully-restores FMRP levels, but also normalizes a hyperexcitable phenotype [29], as well as rescues synaptic scaling deficits [13]. Moreover, demethylation of the extended CGG could also restore *FMR1* levels and attenuate increased spontaneous activity in FXS iPSC-derived neurons [29]. While it has been demonstrated that near complete restoration of *FMR1* levels was able to normalize a hyperexcitable phenotype, there has not yet been a systematic assessment to determine if partial restoration is sufficient to correct this phenotype.

In this study, we utilize two different isogenic iPSC pairs to confirm that the absence of FMRP leads to neuronal hyperactivity using multielectrode arrays (MEAs). We used orthogonal gene expression technologies to titrate the levels of FMRP expression in excitatory human neurons. We then determined the levels of FMRP necessary to correct this phenotype by two separate means: first, the proportion of FMRP positive neurons needed throughout the culture, and second, the total amount of FMRP required as a percentage of control levels. Our data demonstrates that partial restoration of FMRP is sufficient to normalize hyperactivity in FXS iPSC-derived neurons.

## Materials and Methods

### iPSC culture

WT isogenic control and FXS iPSC lines used in this study are listed in supplemental table S1. iPSCs were cultured on Geltrex (Thermo Fisher Scientific, A1413302) coated flasks in StemFlex™ medium (Thermo Fisher Scientific, A3349401) and fed with fresh media every other day. Cells were passaged using ReLeSR™ (STEMCELL Technology, 05873) and tested for mycoplasma and karyotypic abnormalties. iPSCs were cryopreserved in CryoStem™ hPSC Freezing Medium (Biological industries, 05-710-1E). To recover the iPSCs, vials were thawed in a 37°C water bath for 2 min until a small piece of ice was left in the vial. 1 mL thawing media (StemFlex with 1x RevitaCell (Thermo Fisher Scientific, A2644501)) was added dropwise, then transferred to a 50mL conical containing 10mL thawing media. Cryovials were rinsed once with 1 mL thaw media. Cells were spun at 200 g for 5 min at room temperature, and the supernatant aspirated. Cells were resuspended in 10 mL thawing media and plated.

### FMR1 KO isogenic iPSC engineering

To generate FMR1 KO iPSCs, Epi49 WT iPSCs (Thermo Fisher Scientific, A18945) were cultured in Geltrex coated T75 flask. The day before electroporation, cells were fed with fresh Stemflex™ medium with 1X RevitaCell supplements. Cells were dissociated with 5 ml Accutase™ cell dissociation reagent (STEMCELL technology, 07920). After washing once with PBS, cells were resuspended in Resuspension buffer R (Neon™ Transfection System 100μL Kit, Invitrogen, 10431915) to a final cell density ∼ 10^8^/mL. Transfer 60 ug Cas9-2A-GFP-FMR1 gRNA plasmid (with the gRNA sequence 5’-TATTATAACCTACAGGAGGT-3’ targeting against exon 3) into the dissociated iPSCs and electroporate with the following program: Pulse voltage 1,100V; Pulse width 30ms; Pulse number 1; Cell density at 1 × 10^8^ cells/mL. After electroporation, cells were plated into a Geltrex coated T75 flask using Stemflex™ medium with 1 X RevitaCell supplements. On day 3 post electroporation, cells were dissociated with Accutase for FACS sorting to enrich the GFP+ population, and re-plated onto Geltrex coated 10cm Petri dish at ∼ 5,000 cells/plate. 1X RevitaCell was supplemented in the Stemflex medium to enhance the cell viability. iPSC cell colonies were hand-picked and expanded in Stemflex medium from 96 wells to 12 wells, and further expanded for cryopreservation. Two primers HW15 (5’-actatattgccgttatgtcccactcag-3’) and HW16 (5’-taagatgagttagtcaaaagcacgtgtc-3’) were used to generate the PCR amplicon that flanks the gRNA target region in exon 3 of *FMR1*. Based on sequencing results, clone A2 has homozygous frameshift mutation leading to a premature stop codon for FMRP protein with the following truncated peptide sequence MEELVVEVRGSNGAFYKAFVKDVHEDSITVAFENNWQPDRQIPFHDVRFRL*. (Figure 6A)

### Neuronal differentiation of iPSC

NGN2 neurons were derived from human WT and FXS iPSCs according to a modified method described by Zhang et al. (Zhang et al, Neuron, 2013). Briefly, WT and FXS iPS cells were cultured on Geltrex in StemFlex™ medium, and dissociated with TrypLE™ (Thermo Fisher, 12605010) as single cells. Cells were infected with pTet-O-NGN2-2A-Puro virus (1 × 10^9^ IFU/mL) in suspension at MOI ∼ 5 and plated in Stemflex with 1X RevitaCell. Infected cells were cultured and expanded on Geltrex in StemFlex media. On differentiation day 0, cells were dissociated with TrypLE™ and replated in StemFlex with RevitaCell and 2 μg/mL Doxycycline (Fisher Scientific, BP2653-1) to induce TetO gene expression. Media was changed 24hrs post induction to DMEM/F12 (Thermo Fisher, 11320033) with 1X N2 supplement (Thermo Fisher, 17502048), 1X Glutamax (Thermo Fisher, 35050061), 0.3% D-(+)Glucose (Sigma Aldrich, G8769), 2 μg/mL Doxycycline and 3.34 μg/mL Puromycin (Thermo Fisher, A1113808) to start selection. 48 h post-induction, the media was changed to differentiation media containing 1X B27 supplement (ThermoFisher, 17504044). Neurons were dissociated on day 3 with room temperature StemPro™ Accutase™ Cell Dissociation Reagent (Thermo Fisher, A1110501) and re-plated into PDL and Laminin (ThermoFisher, 23017-015) coated plates in NBM media (Neurobasal™ Media (ThermoFisher, 21103049) with 1X MEM NEAA (Thermo Fisher, 11140050), 1X Glutamax, 0.3% D-(+)-Glucose, 2 μg/mL Doxycycline, 3.34 μg/mL Puromycin, 1X B27 supplement, and 10 ng/mL rhBDNF (R&D Systems, 248-BDB/CF)) and rhGDNF (R&D Systems, 212-GD)). The day following plating, fresh NBM was added to the cultures. Half media changes were conducted weekly.

### qRT-PCR

Cells were lysed using Real-Time ready cell lysis buffer (Roche Life Science, 7248431001) with 1:80 diluted RNAse inhibitor (Sigma, 3335402001) and DNAseI (ThermoFisher Scientific, AM2222). 96-well plates were lysed by aspirating the media the cells and adding 50 μL lysis buffer per well. The plates were shaken at 1250 RPM for 25 min. The lysate was mixed with a pipette 10 times and checked for lysis under the microscope. Plates were frozen for at least 15 min at −80°C. Lysates were thawed on ice and diluted 1:5 with nuclease-free water. 2.25 μL of diluted total lysate was used in a PreAmp reaction with the FMR1-FAM and POP4-ABY TaqMan assays (5 μL PreAmp Master Mix (Applied Biosystems 4488593), 2.5 μL 0.2X diluted FMR1 and POP4 assays, 0.25 μL 40X TaqMan RT enzyme mix (Applied Biosystems A36107C), 2.25 μL lysate). The preamp reaction was run for 12 cycles in an Applied Biosystems QS7 machine. PreAmp reactions were diluted 1:5 and 2 μL used in a qPCR reaction with the same two TaqMan assays (5uL TaqMan Fast Advanced Master Mix (Applied Biosystems 4444557), 0.5 μL 20X FMR1-FAM TaqMan assay, 0.5 μL 20x POP4-ABY TaqMan assay, 2 μL Nuclease-Free water) and run on a QuantStudio 7 (Thermo
Fisher Scientific).

### Immunofluorescence and imaging analysis

Neurons were fixed with 4% paraformaldehyde (PFA) for 10 min at room temperature. Cells were permeabilized with PBST (1X PBS solution with 0.5% Triton X-100) before blocking with 5% BSA in PBST. Cells were then incubated with appropriately diluted primary antibodies in PBST with 1% BSA for 1 h at room temperature or overnight at 4°C. After washing with PBST for 3 × 10 min at room temperature, cells were incubated with fluorophore labeled secondary antibodies and DAPI. Cells were washed three times with PBST at room temperature and subjected to imaging using CellInsight CX7 High-Content Screening (HCS) Platform (Thermo Fisher Scientific). Data were analyzed using Cellomics software and plotted with Prism GraphPad for statistical analysis. For imaging-based quantification, unless otherwise specified, >10 representative images were quantified, and data were plotted as mean±SD. Dilution for antibodies used in this study is listed in supplemental table 3.

### MEA and data analysis

48 well multielectrode array plates containing an array 16 embedded gold electrodes were used for all MEA experiments (Cat. #M768-KAP; Axion BioSystems, Atlanta, GA). The day before cell plating, MEA wells were coated with a 0.07% polyethylenimine (PEI) solution (Sigma, St. Louis, MO) for 2 h in a 37°C cell-culture incubator. Plates were rinsed with sterile, deionized water four times and air dried in a biological safety cabinet overnight. Neurons were cultured in BrainPhys medium (STEMCELL Technology) supplemented with 1X N-2 Supplement (ThermoFisher), 1 μg/mL Laminin (Sigma-Aldrich), Neural Supplement B (Cellular Dynamics, M1029), Nervous System Supplement (Cellular Dynamics, M1031), and 1X Penicillin-Streptomycin (ThermoFisher). On the day of plating, a 10µL drop of complete BrainPhys media containing 80 μg/mL Laminin was placed directly in the center of each well over the MEA electrodes. The plates were then incubated for at least 1 h at 37°C. A 10µL drop of cell suspension was then plating directly into the 10µL drop of complete BrainPhys media containing 80 μg/mL Laminin, for a final volume of 20µL and final density of 75,000 neurons per well. The cells were allowed to attach to the substrate in a cell-culture incubator for 1 h, and then 500 μL culture medium was added to the wells. Cells were grown at 37°C in a humidified 5% CO_2_/95% O_2_ incubator. To maintain the culture, a 50% medium change with fresh 37°C culture medium was performed every 3–4 days.

Spontaneous neuronal activity, including burst rate and firing rate, were recorded every day using the Axion Maestro MEA platform (Axion Biosystems). Recording was performed every day before or after dCas9-Tet1/CGG sgRNA, *FMR1* mRNA and *FMR1* ASO treatments. Data were analyzed using a custom MATLAB program [30] and presented as mean ± SD, and are considered statistically significant when p < 0.05 by paired Student’s T-Test or one-way ANOVA, where appropriate.

### FMR1 mRNA Transfection

*FMR1* mRNA was designed with full substitution of pseudo-U and ARCA capped to enhance the stability and translation efficiency. The mRNA was synthesized by TriLink BioTechnologies (L-6009) and purified with Silica membrane with enzymatical polyadenylation and Dnase treatment. mRNA transfection was performed using Lipofectamine™ MessengerMAX™ Transfection Reagent (ThermoFisher Scientific, LMRNA003) according to manufacturer’s instruction. Briefly for transfection of each well, various amounts of mRNAs were diluted in 15 μl OptiMEM and combine with prediluted 15 μl 2X Lipofectamine MessengerMAX reagent (0.5 μl/well) in equal volume of OptiMEM. Transfectant were fully mixed by vortexing and incubation at RT for 5 min. 30 μl transfection cocktail were added into each well of the neuronal culture for transfection. Change 75% of the media 16 h after transfection. A synthetic CleanCap™ EGFP mRNA (TriLink, L-7201) was used as a transfection positive control in this study.

### dCas9-Tet1 construct and lentiviral transduction

The Fuw-dCas9-Tet1CD-P2A-BFP (Addgene, 108245), Fuw-dCas9-dTet1CD-P2A-BFP (Addgene, 108246) and CGG sgRNA-mCherry co-expression plasmids were gifts from Rudolf Jaenisch lab at Whitehead Institute, MIT, Cambridge, MA. The *FMR1* CGG sgRNA expression construct was cloned by inserting annealed oligos into modified pgRNA plasmid (Addgene plasmid: 44248) with AarI site. To package lentivirus, a synthetic gene block encoding the bacteriophage AcrIIA4 purchased from IDT was cloned into a modified FUW vector [29]. All constructs were sequenced before lentiviral packaging and production at 1 × 10^9^ IFU/mL by Alstem LLC. For lentiviral transduction, neurons were infected with combination of dCas9-Tet1CD/CGG sgRNA and dCas9-dTet1CD/CGG sgRNA at various MOI, media were changed 16 h post transduction.

### FMR1 ASO treatment

Two *FMR1* ASOs were designed and synthesized by EXIQON (FMR1 ASO #1: Product sequence: ACAGATCTTAGACGTT, Cat. No. LG00199842; FMR1 ASO #2: Product sequence: GGAATAAGAATTACGG, Cat. No. LG00199843). For *FMR1* ASO titration, various amount of *FMR1* ASOs (10nM to 1μM) were added into WT neurons cultured on PDL/Laminin coated plate for imaging analysis or PEI/Laminin coated MEA plates for electrophysiological analysis.

## Results

### Quantification of FMRP-expression in isogenic pair of iPSC-derived neurons

In this study, we utilized an isogenic pair of iPSCs in which the expression level of FMRP was the differentiating variable. This pair was generated by identifying clonal lines following CRISPR-mediated removal of the CGG-repeat region from the *FMR1* gene of a human FXS iPSC line harboring a >450 CGG repeat expansion [27]. We have previously confirmed both the lack of *FMR1* expression and increased spontaneous activity in these FXS iPSC-derived cortical excitatory neurons as compared to CRISPR-edited isogenic control [29]. Figure 1A shows representative immunofluorescent images from both the FXS iPSC-derived neurons (SW_FXS) and isogenic control neurons (SW_C1_2) demonstrating robust FMRP staining in cell bodies of control neurons compared to no FMRP staining in the FXS iPSC-derived neurons. Figure 1A also demonstrates that both cell lines display multiple neuronal markers such as β-Tubulin, MAP2 and NeuN. FMRP levels are quantified in Figure 1B by the percentage of FMRP+ neurons, with close to 80% of C1_2 neurons positive for FMRP, and around 1% of FXS neurons positive for FMRP (C1_2: 79.2±15.4%, n=18 wells; FXS: 1.2±1.3%, n=24 wells).

**Figure 1.**
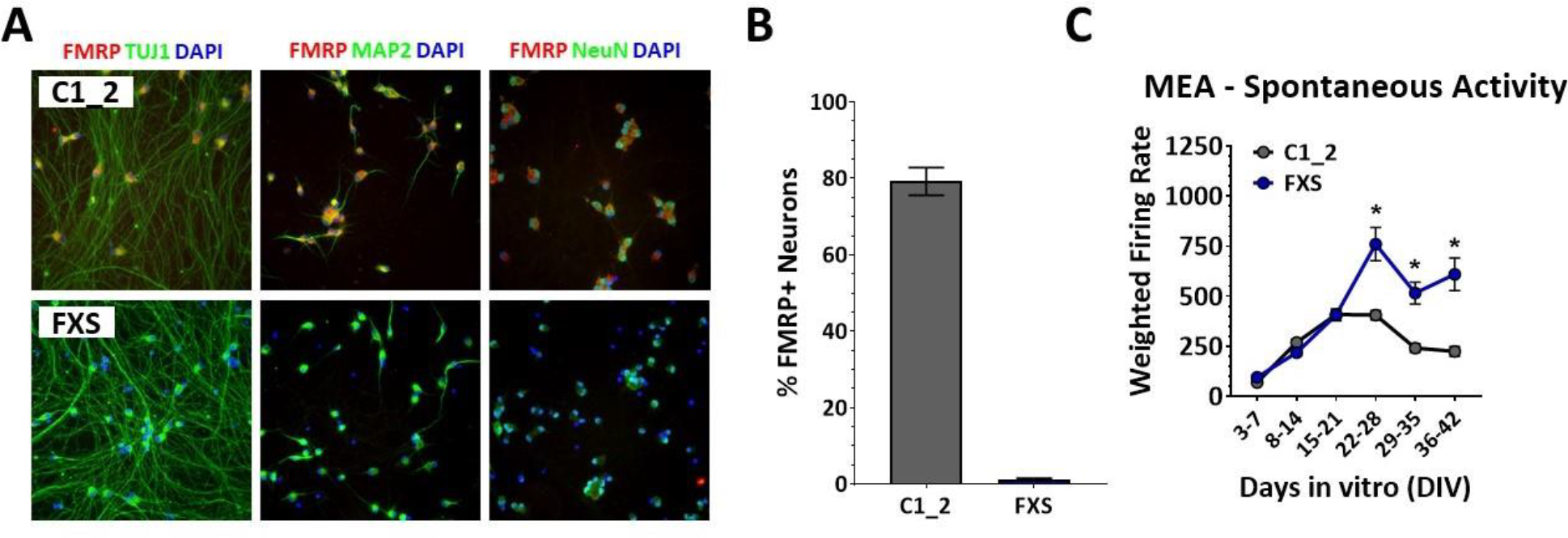
iPSC-derived neurons from a Fragile X patient do not express FMRP and show increased spontaneous activity. **A**. Immunofluorescent images from FXS iPSC-derived neurons (FXS) and a CRIPSR/Cas9-corrected isogenic control (C1_2) stained for nuclei (blue) FMRP (red) and neuronal markers (green) demonstrate no FMRP staining FXS neurons (lower 3 panels), compared to robust FMRP staining in the cell soma for corrected C1_2 neurons (upper 3 panels). The neuronal markers (green) used are β-Tubulin (first column), MAP2 (second column) and NeuN (third column). **B**. Quantification of FMRP positive neurons for both FXS and C1_2 neurons demonstrates that the majority of corrected C1_2 neurons are positive for FMRP, whereas FXS neurons are not. **C**. Increased spontaneous activity as measures by multi-well MEAs shows that increased FXS neuronal activity begins to manifest after 3 weeks in culture.

### Absence of FMRP results in increased spontaneous activity

To confirm published work demonstrating increased spontaneous activity in excitatory FXS iPSC-derived neurons, we compared the levels of spontaneous electrical activity for both isogenic lines using multi-well MEA plates. The level of spontaneous activity was quantified using an average weighted firing rate per well, which was calculated as a product of the spontaneous spikes per minute and the number of active electrodes in the well. Figure 1C shows the spontaneous activity for all three isogenic pairs of iPSC-derived neurons over the course of seven weeks. These data demonstrate that neurons lacking FMRP expression begin to show increased spontaneous activity after 3 weeks which is maintained throughout the remaining time in culture. Cumulative data from the fourth week shows a significantly higher weighted firing rate for FXS iPSC-derived neurons when compared to the corrected isogenic control (FXS: 760.5±918.0, n=120 wells, C1_2: 406.0±254.6, n=118 wells, *p<0.01, two-way ANOVA, Sidak’s multiple comparison test).

### Mixed neuronal culture determines the level of FMRP mosaicism for functional restoration

We next sought to determine how many FMRP positive neurons would be needed to normalize the increased spontaneous activity seen in FXS neuronal cultures. To do this, we generated mixed cultures of FMRP-expressing (C1_2) and FMRP-non-expressing (FXS) neurons by titrating specific ratios of C1_2 neurons into FXS cultures. This paradigm allowed for the generation of a panel of FMRP mixed cultures that mimic neuronal environments in FXS mosaic male patients and female carriers. Figure 2A shows representative images of from 100% C1_2 and 100% FXS neuronal cultures, along with FXS cultures in which 1% and 20% C1_2 neurons have been mixed with 99% and 80% FXS neurons respectively. The percentage of C1_2 neurons for each of these mixed cultures is quantified by ICC in Figure 2B, showing increasing amounts of FMRP positive neurons as the ratio of C1_2 to FXS neurons is increased. When the level of spontaneous activity was measured for the different mixed-ratio cultures, a significant increase in spontaneous activity was seen in 100% FXS cultures as compared to 100% C1_2 (100% C1_2: 1.00±0.97, n=21 wells; 100% FXS: 5.38±6.01, n=30 wells; *p<0.01, one-way ANOVA, Dunnett’s multiple comparison test). As C1_2 neurons were titrated into FXS cultures, only FXS cultures in which 20% or greater C1_2 neurons are present showed significant decreases in spontaneous activity when compared to the 100% FXS cultures. Cultures which had 20% or greater C1_2 neurons demonstrated spontaneous activity levels comparable to 100% C1_2 levels, suggesting that at least 20% FMRP-expressing neurons are needed to normalize increased spontaneous activity seen in FXS neuronal cultures (100% C1_2: 1.00±0.97, n=21 wells; 20% C1_2: 1.25±1.17, n=28 wells, 50% C1_2: 1.53±1.30, n=26 wells, p>0.9, one-way ANOVA, Dunnett’s multiple comparison test).

**Figure 2.**
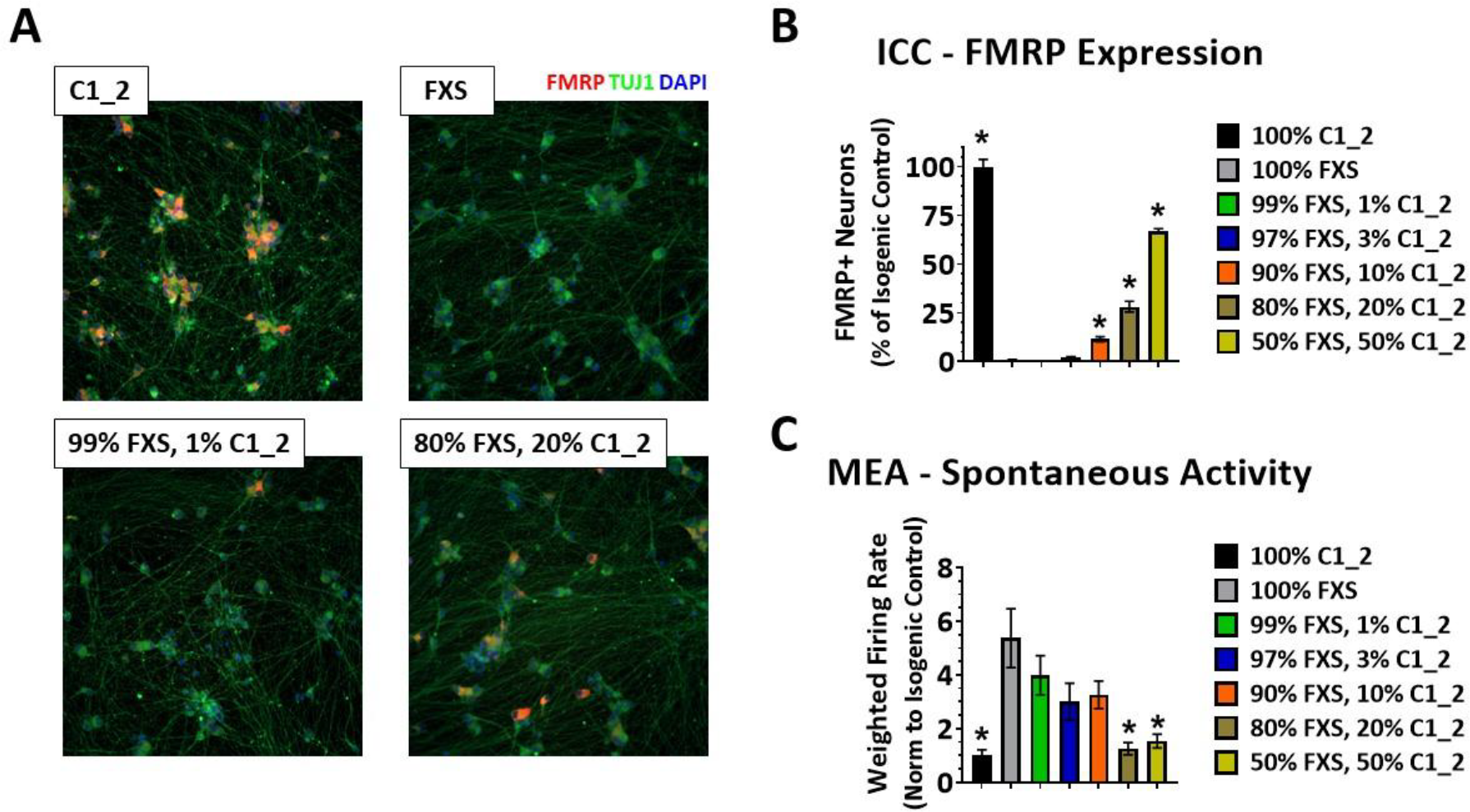
Mixing FMRP positive and FMRP negative neurons results in a FMRP mosaic culture. **A**. Immunofluorescent images from C1_2 neurons (upper left panel) and FXS neurons (upper right panel) in which 1% (lower left panel) and 20% FMRP-positive neurons (lower right panel) have been mixed. Blue – nuclei, green – β-Tubulin, red – FMRP. **B**. Quantification of FMRP-positive neurons in the different FMRP-mosaic cultures shows increasing FMRP-positive neurons with increasing percentages of C1_2 neurons mixed into FXS cultures. Data normalized to C1_2 cultures. **C**. Quantification of spontaneous activity recorded on MEAs three weeks post plating demonstrates reduced neuronal activity in FXS neuronal cultures in which greater than 20% FMRP-expressing neurons are present.

### Transient transfection of WT FMR1 mRNA results in mosaic FMRP expression pattern to rescue FXS functional phenotype

To further explore the levels of FMRP-positive neurons needed to normalize FXS neuronal hyperactivity, we transfected varying amounts of WT human *FMR1* mRNA into FXS cultures. Due to the stochastic nature of mRNA uptake during transfection, this experimental paradigm produced a FMRP mixed culture similar to that seen with cell mixing. Representative images of C1_2, FXS and FXS plus 8 or 16ng of FMR1 mRNA are shown in Figure 3A. The lower two panels demonstrate the stochastic nature of FMRP expression 24hr post transfection in which neurons are either FMRP positive or FMRP negative, with the number of FMRP-expressing neurons dependent upon the amount of transfected *FMR1* mRNA. The percentage of FMRP-expressing neurons is quantified in Figure 3B, showing that as the amount of *FMR1* mRNA is increased, the percentage of FMRP+ neurons also increases, with a maximal level of around 25% FMRP positive neurons achieved 24h after transfecting. This FMRP expression at 24h represented the peak expression levels, with levels beginning to decrease 48-72h later. Due to the transient nature of FMRP expression following *FMR1* mRNA transfection, the greatest effect on spontaneous activity was seen 24-32h post transfection (Figure 3C). Here, the data from 3 recording sessions at 24h, 28h and 32h post-transfection were averaged and normalized to a pre-transfection baseline recording. From this data, a significant difference in spontaneous activity between C1_2 and FXS is seen during this post-transfection time period. In addition, only FXS cultures that were transfected with 24ng of *FMR1* mRNA demonstrated a significant reduction in spontaneous activity compared to the mock treated FXS neuronal cultures (FXS: 2.5±2.2, n=123; FXS + 24ng *FMR1* mRNA: 1.6±2.8, n=153, *p<0.05, one-way ANOVA, Dunnett’s multiple comparison test). This level of FMR1 mRNA corresponded to 24.6±5.5% FMRP-positive neurons (n=6 wells). These data support the conclusion that greater than 20% FMRP-expressing neurons normalize the increased spontaneous activity seen in FXS cortical neuronal cultures.

**Figure 3.**
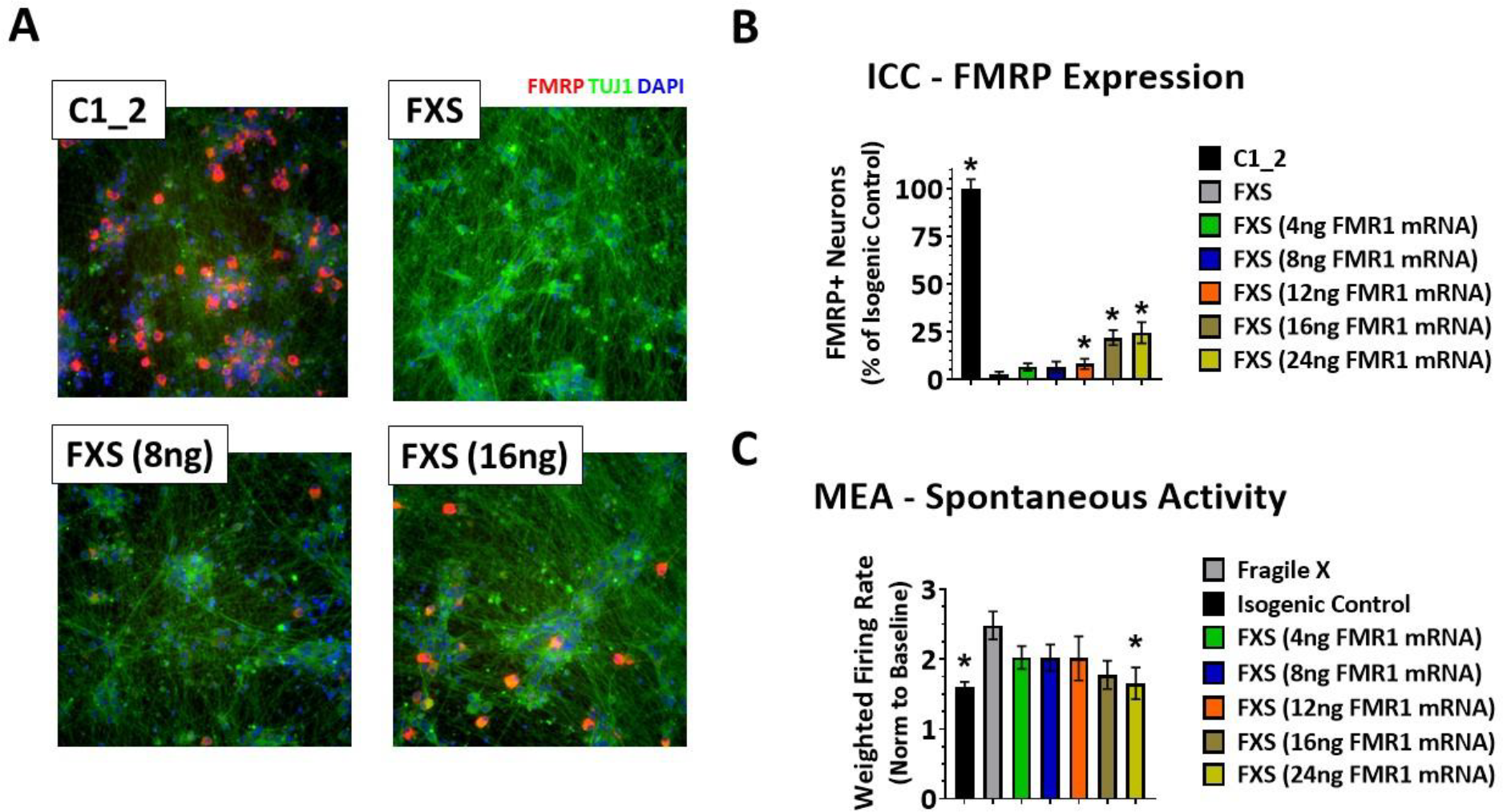
Transfection of FMR1 mRNA into FXS cultures results in a transient FMRP mosaic culture. **A**. Immunofluorescent images from C1_2 neurons (upper left panel) and FXS neurons (upper right panel) in which 8ng FMR1 mRNA (lower left panel) and 16ng FMR1 mRNA (lower right panel) have been transiently transfected. Blue – nuclei, green – β-Tubulin, red – FMRP. **B**. Quantification of FMRP-positive neurons in cultures 24hr following transfection with varying amounts of FMR1 mRNA shows an increasing number of FMRP-positive neurons with increasing amounts of FMR1 mRNA. Data normalized to C1_2 cultures. **C**. Quantification of spontaneous activity, normalized to a pre-transfection baseline, recorded on MEAs beginning 24hr post transfection of varying amounts of FMR1 mRNA, shows reduced neuronal activity in FXS neuronal cultures in which greater than 25% FMRP-expressing neurons are present

### Titration of FMR1 ASO leads to varying reduced levels of FMRP reduction in WT neurons results in increased spontaneous activity

After establishing that greater than 20% FMRP-expressing neurons were needed to normalize the increased spontaneous activity seen in FXS cultures, we next sought to determine the overall expression levels of FMRP, rather than the number of highly-expressing FMRP neurons, that are required to normalize FXS spontaneous activity. To achieve this, we first attempted to uniformly lower *FMR1* levels across the culture using antisense oligonucleotides (ASOs) targeted to the *FMR1* mRNA in C1_2 neurons (Figure 4). Figure 4A demonstrates the level of both *FMR1* and FMRP following a 2wk treatment with two different *FMR1*-targeting ASOs. Here, a robust knockdown of *FMR1* is seen following treatment with both ASOs at 1 µM (ASO #1: 28.8±1.8% of control, n= 3; ASO #2: 26.6±2.8% of control, n= 3). An even greater reduction in FMRP was observed following ASO treatment, with ASO #1 producing around a 93% reduction in FMRP and ASO #2 producing around a 96% reduction (ASO #1: 7.0±0.5% of control, n= 3; ASO #2: 4.4±2.0% of control, n= 3). When the spontaneous activity of C1_2 cultures was recorded 2 weeks post-ASO treatment, only cultures treated with 1µM ASO #2 showed a significant increase in spontaneous activity as compared to C1_2 cultures treated with a non-targeting control ASO. Cultures treated with 1µM ASO #1 did not show a significant increase in spontaneous activity, suggesting that 7% FMRP expression within the culture was sufficient to prevent the increased spontaneous activity, whereas the 4% FMRP expression produced with 1µM ASO #2 was a great enough reduction to induce increase spontaneous activity (Figure 4B; C1_2 + 1µM Control ASO: 57.0±50.0, n=21; C1_2 + 1µM ASO #1: 90.8±92.5, n=9; C1_2 + 1µM ASO #2: 241.4±188.8, n=12; *p<0.05, one-way ANOVA, Dunnett’s multiple comparison test). These data suggest that while greater than 20% FMRP-expressing neurons are needed to normalize increased spontaneous activity in FMRP mosaic cultures, a lower level of FMRP expression across all neurons in a culture may be enough to normalize increased spontaneous activity due to the absence of FMRP.

**Figure 4.**
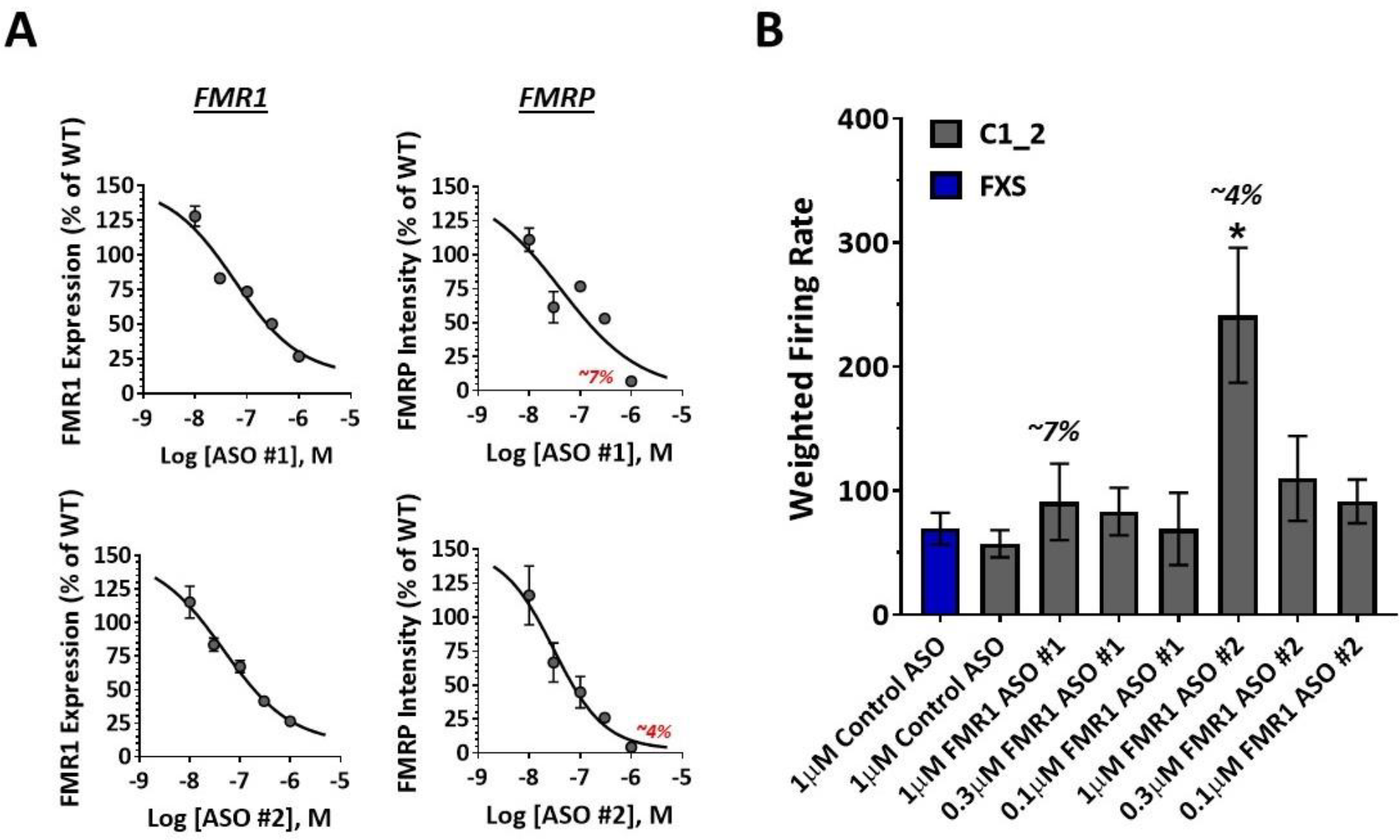
ASO-mediated reduction of FMR1 and FMRP in FMRP-expression neurons results in increased spontaneous activity. **A**. Concentration response curves for relative FMR1 expression (first column) and FMRP expression (second column) following a two-week treatment with varying concentrations of two different ASOs shows a robust, concentration-dependent knockdown of both FMR1 and FMRP. **B**. Quantification of spontaneous activity recorded on MEAs two weeks post-ASO treatment demonstrates increased activity only in C1_2 cultures in which greater than 95% FMRP knockdown is achieved, as denoted by the bar labeled 4%.

### Titration of dCas9-Tet1/CGG sgRNA leads to increasing levels of FMRP in FXS neurons and functional rescue

In addition to knocking down FMRP in FMRP-expressing C1_2 neurons, we also attempted to reactivate FMRP in FXS neurons. To this end, we utilized a catalytically-inactive Cas9 protein fused to the methylcytosine dioxygenase Tet1 as targeted DNA demethylation editing machinery. We have previously shown that this construct, when targeted to the CGG-repeat region of the FMR1 gene, can demethylate this hypermethylated region and reactivate FMR1, resulting in FMRP expression in multiple FXS iPSCs and neurons [29]. Here, we transduced iPSC-derived FXS neurons with low titers of the dCas9-Tet1 construct to produce partial reactivation of FMR1 (Figure 5). Figure 5A depicts representative images from FXS, C1_2 and FXS neurons transduced with 1 MOI of the dCas9-Tet1 construct. These images show no FMRP expression in the FXS neurons, robust FMRP expression in the C1_2 neurons, and low expression of FMRP in the majority of FXS neurons following transduction with dCas9-Tet1/CGG gRNA lentiviruses at 1 MOI. The quantification of FMRP intensity, normalized to C1_2 is shown in Figure 5B. Here, FXS neurons show less than 1% FMRP intensity as compared to C1_2 neurons (C1_2: 100±14.9%, n=6 wells; FXS:0.9±0.7%, n=6 wells). A non-significant increase in FMRP intensity is seen in both FXS neurons transduced with a high MOI of a catalytically-inactive Tet1 (dCas9-dTet1) control and a low MOI of the active dCas9-Tet1 construct (5 MOI dCas9-dTet1: 3.4±3.8%, n=6 wells; p=0.4, one-way ANOVA, Dunnett’s multiple comparison test; 0.5 MOI dCas9-Tet1: 5.4±2.5%, n=6 wells; p=0.06, one-way ANOVA, Dunnett’s multiple comparison test). A statistically significant increase in FMRP intensity of around 7% was seen in FXS neurons transduced with 1 MOI of the active dCas9-Tet1 as compared to non-transduced FXS control neurons (FXS:0.9±0.7%, n=6 wells; 1 MOI dCas9-Tet1: 7.1±4.4%, n=6 wells; *p<0.01, one-way ANOVA, Dunnett’s multiple comparison test). When spontaneous activity levels were measured following transfection, both control FXS neurons and FXS neurons transduced with the control dCas9-dTet1 construct showed statistically-significant increases in spontaneous activity as compared to FMRP-expression C1_2 neurons, whereas FXS neurons transduced with 0.5 and 1 MOI of the dCas9-Tet1 construct did not (Figure 5C; C1_2: 65.6±174.5, n=479 electrodes, FXS: 121.3±272.7, n=479 electrodes; 5 MOI dCas9-dTet1: 144.4±308.9, n=463 electrodes; 0.5 MOI dCas9-Tet1: 97.0±213.4, n=451 electrodes; 1 MOI dCas9-Tet1: 82.7±210.8, n=449 electrodes; *p<0.01, one-way ANOVA, Dunnett’s multiple comparison test). These data indicate that >7% FMRP levels is sufficient to normalize increased spontaneous activity in FXS neurons – a similar threshold level identified through our FMR1 ASO experiments in FMRP expressing C1_2 neurons in which greater than 95% FMRP reduction (or <5% FMRP levels) was needed to induce an increased spontaneous activity phenotype.

**Figure 5.**
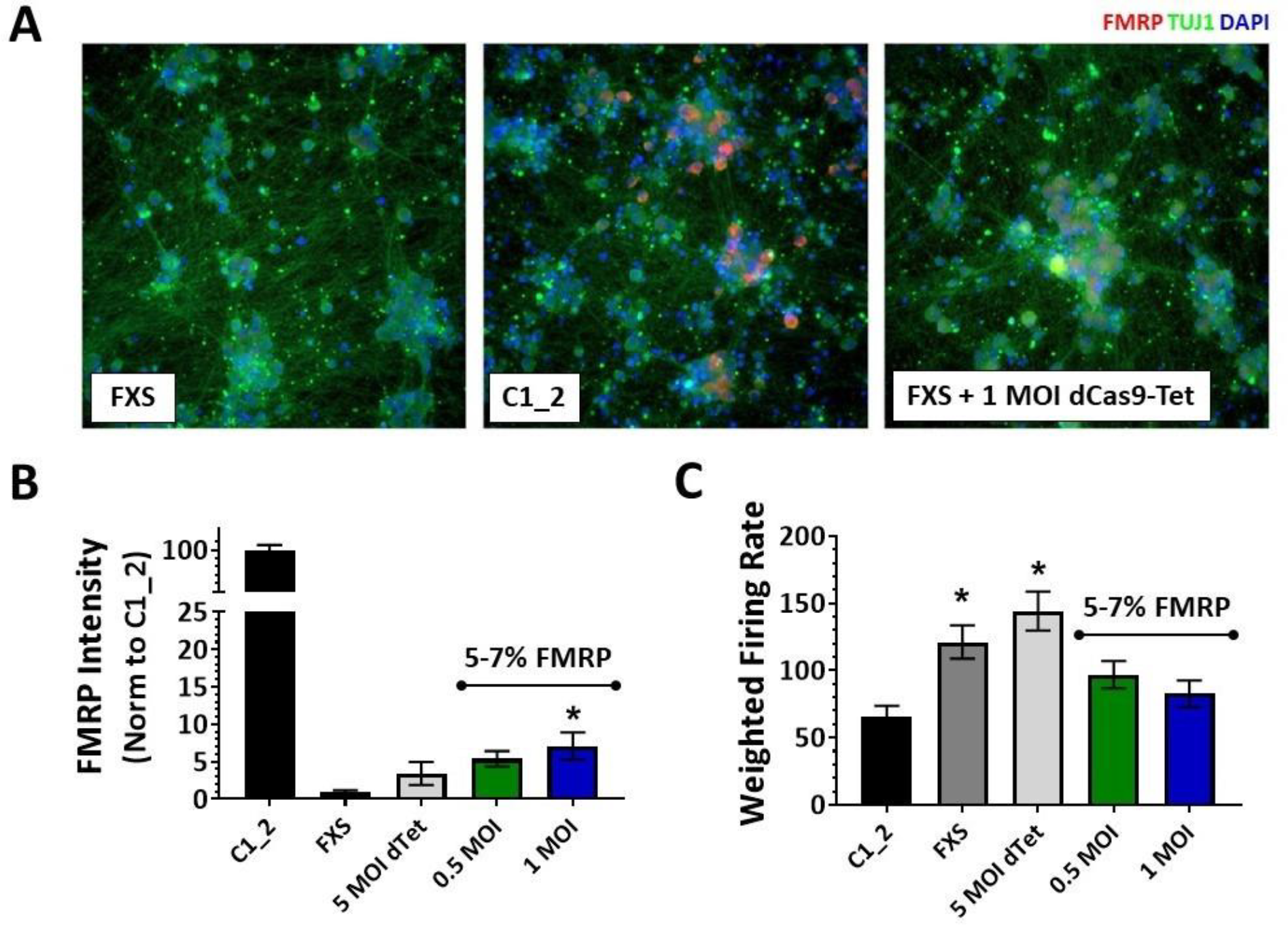
Partial reactivation of FMRP using a dCas9-Tet1 construct reduces increased spontaneous activity. **A**. Immunofluorescent images from FXS neurons (left panel), C1_2 neurons (middle panel), and FXS neurons 2 weeks post-transduction of 1 MOI dCas9-Tet1. Blue – nuclei, green – β-Tubulin, red – FMRP. **B**. Quantification of FMRP intensity for C1_2 neurons and FXS neurons transfected with 5 MOI dCas9-dTet1, 0.5 MOI or 1 MOI dCas9-Tet1 shows increase FMRP intensity with 1 MOI dCas9-Tet1. Data normalized to C1_2. **C**. Quantification of spontaneous activity recorded on MEAs two weeks post transduction demonstrates reduced spontaneous activity from FXS cultures with greater than 5% of C1_2 FMRP expression.

### Confirmation of neuronal hyperactivity phenotype in genetically engineered FMRP KO neurons

Finally, we utilized a commercially-available healthy, normal iPSC line to create a second pair of isogenic iPSC-derived neurons that either did or did not have detectable levels of FMRP. Here, a *FMR1* knockout line (FMRP KO) was created by inserting a premature stop codon in the third exon of the *FMR1* gene, which resulted in normal levels of truncated FMR1 mRNA but no expression of FMRP (Figure 6A). Representative immunofluorescent images for this pair of isogenic iPSC-derived neurons are shown in Figure 6B. In addition, electrical activity recorded from these lines demonstrated an increased spontaneous activity phenotype in FMRP KO neurons that developed after 3 weeks in culture as compared to wild-type (WT), a similar timeframe seen with the first pair of isogenic neurons we profiled (Figure 6C; FMRP KO: 418.2±427.9, n=21 wells, WT: 120.1±123.4, n=15 wells, *p<0.01, two-way ANOVA, Sidak’s multiple comparison test). Moreover, similar results were obtained when we performed a cell mixing experiment in which WT neurons were titrated into a culture of FMRP KO cells. These results are quantified in Figure 6D showing increasing amounts of FMRP-positive neurons as an increasing percentage of FMRP-expression control neurons are titrated into FMRP KO cultures. A statistically-significant increase in FMRP positive neurons is seen in both 80% FMRP KO / 20% Control (36.6±18.3%, n=6 wells; *p<0.01, one-way ANOVA, Dunnett’s multiple comparison test) and 50% FMRP KO / 50% Control (53.2±24.8%, n=6 wells; *p<0.01, one-way ANOVA, Dunnett’s multiple comparison test). When spontaneous activity levels were recorded from these cultures, only cultures with 20% or greater FMRP-expressing neurons show a significant reduction in spontaneous activity levels as compared to the 100% FMRP KO neurons (Figure 6E; 100% control: 1.0±0.6, n=32 wells; 100% FMRP KO: 1.8±1.1, n=31 wells; 80% FMRP KO / 20% control: 0.9±0.3, n=12 wells; 50% FMRP KO / 50% control: 1.0±0.5, n=12 wells; *p<0.05, one-way ANOVA, Dunnett’s multiple comparison test). These data support an FMRP-dependent increased spontaneous activity phenotype, as well as confirm a threshold level of >20% FMRP-expressing neurons needed in a FMRP mosaic culture to normalize increased spontaneous neuronal activity.

**Figure 6.**
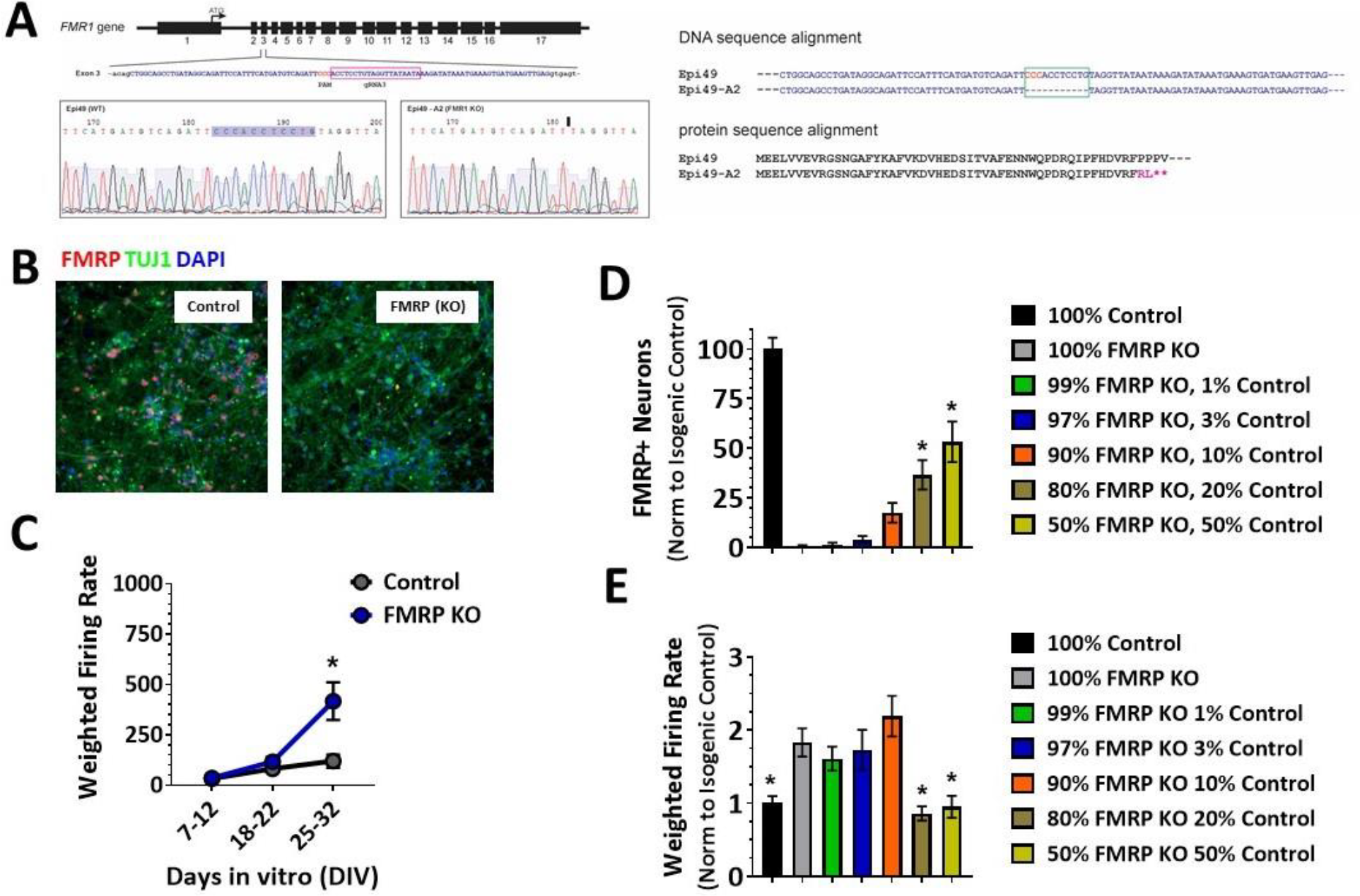
FMRP knockout neurons shown increased spontaneous activity that is normalized via partial addition of FMRP-expression neurons. **A**. Generation of *FMR1* KO iPSC by CRISPR/Cas9 mediated genetic editing from a WT iPSC line (Epi49). Left, a gRNA was designed to target the exon 3 of *FMR1* locus from 3’ to 5’ ends. Amplicon sequence identified Clone A2 with homozygous deletion in the gRNA targeting region. The deleted sequence and the new ligation joint were highlighted in the sequence profiles of WT and the A2 clone. Right, DNA and protein sequence alignments of the WT and *FMR1* KO iPSC indicate that the *FMR1* KO clone produces an N-terminal truncated polypeptide. Asterisks denote stop codons. **B**. Immunofluorescent images from control WT iPSC-derived neurons (left panel) and neurons in which FMRP has been knocked out. Blue – nuclei, green – β-Tubulin, red – FMRP. **C**. Increased spontaneous activity as measures by multi-well MEAs shows that increased neuronal activity in FMRP KO neurons begins to manifest after 3 weeks in culture. **D**. Quantification of FMRP-positive neurons in the different FMRP-mosaic cultures shows increasing FMRP-positive neurons with increasing percentages of WT neurons mixed into FMRP KO cultures. Data normalized to control cultures. **E**. Quantification of spontaneous activity recorded on MEAs three weeks post plating demonstrates reduced neuronal activity in FMRP KO neuronal cultures in which greater than 20% FMRP-expressing neurons are present.

## Discussion

In this study, we have confirmed previous reports of increased spontaneous activity in FXS patient-derived excitatory neurons differentiated from iPSCs using two isogenic pairs of cell lines in which the presence or absence of FMRP is the dependent variable. In addition, we have demonstrated that this increased neuronal activity can be normalized through partial restoration of FMRP-expressing neurons within the neuronal network. Our results suggest that the number of FMRP-expressing neurons within mosaic cultures in which neurons are either expressing FMRP or not, needs to be greater than 20% in order to normalize the increased spontaneous activity. In addition, our results suggest that the overall level of FMRP expression throughout all neurons within the culture needs to be greater than 5% of normal FMRP levels in order to restore the level of spontaneous activity to control levels. These data help to provide threshold levels for potential therapeutics aimed at either fully reactivating *FMR1* in a subset of neurons or partially-reactivating *FMR1* across large neuronal populations.

We utilized two different paradigms to generate FMRP mosaicism in our iPSC-derived neuronal cultures – titrating varying percentages of FMRP-expressing neurons into FXS cultures and transiently transfecting FXS neurons with different amounts of FMR1 mRNA. These FMRP mosaic cultures were composed of different percentages of FMRP positive neurons. Based on these experiments, our data suggest that greater than 20% FMRP positive neurons are needed to normalize increased spontaneous neuronal activity. This number is slightly lower than previously reported percentages of FMRP positive lymphocytes in FXS patients needed for predicting mental retardation. These cut-off levels have been reported to range between 28 and 50% [31–34]. Additionally, a case report has been described in which a FXS patient with 22% FMRP positive leukocytes presented with normal physical features and a high-functioning status, compared to his fully affected brother with a typical FXS presentation and no detectable FMRP [35]. Another study that analyzed FMRP expression in both peripheral blood cells and primary fibroblasts from full mutation and mosaic FXS patients concluded that low, but non-zero expression of FMRP may be sufficient to impact cognitive function [36]. It should be noted that these studies were evaluating FMRP expression in peripheral cells, not in neurons, in which FMRP levels may not correlate. Additionally, there are multiple reports that relate the percentage of FMRP-positive hair follicles – which, like brain tissue, also originate from the ectoderm during embryonic development – to cognitive function, whereas the percentage of FMRP-positive lymphocytes does not [37,38]. These clinical reports support the hypothesis that partial restoration of FMRP may be sufficient to normalize the FXS phenotype, and that low level neuronal FMRP reactivation throughout the brain can positively impact FXS patients. Our study is the first to demonstrate that mixed FMRP cell cultures models can recapitulate the mosaicism observed in some FXS patients, in that similar partial FMRP expression may be sufficient to normalize neuronal activity in the disease.

The use of patient iPSC-derived neurons as a physiologically-relevant model of CNS disease, in conjunction with genome editing techniques to generate isogenic controls, has revolutionized the ability to model the effects of specific disease mutations in a dish [39]. This especially holds promise for therapeutic approaches in FXS that are focused on reactivation of *FMR1* as a disease-modifying strategy. Currently, there are no animal models of FXS that allow for investigating mechanisms of *FMR1* reactivation. The most widely-used model, the *Fmr1* KO mouse, cannot be used to probe potential approaches targeting the root cause of the disease due to the absence of the *Fmr1* gene rather than its inactivation. In addition, attempts to knock-in CGG-repeat expansions in the mouse have not resulted in hypermethylation of the repeat region and silencing of *Fmr1* [40,41]. Therefore, since FXS patient iPSCs retain expanded CGG repeats, promoter hypermethylation and FMR1 silencing after reprogramming [25,26], they represent an attractive disease-relevant model for investigating potential mechanisms for reactivating *FMR1*. Indeed, proof of concept studies showing reactivation of *FMR1* in FXS-derived iPSCs and differentiated neurons have been demonstrated [27,28]. Additionally, several groups are now using FXS patient ESCs and iPSCs to model functional effects in induced FXS human neurons [13,29,42–44] that can be corrected with fully-restored FMRP levels via either CRISPR/Cas9-mediated removal of the CGG repeat region [13,29] or demethylation of the CGG repeat region [29].

The above-mentioned studies have laid the groundwork for demonstrating that FXS functional phenotypes can be corrected *in vitro*. Our study is the first to attempt to understand the minimal levels of FMRP needed to normalize a specific functional phenotype – spontaneous network activity using MEAs. As such, potential small molecule therapeutics targeting *FMR1* reactivation should aim to reach a critical threshold level of FMRP expression in order increase the likelihood of disease modification in patients. Several small molecule therapeutics have failed in the clinic to address various FXS symptoms [24]. We investigated the ability of some these compounds, along with other compounds that have been shown to reactivate *FMR1* in non-neuronal cells [45–47], to increase the expression of *FMR1* in our iPSC-derived patient neurons. Our results however showed that we were unable to demonstrate significant reactivation of *FMR1* following a three-week treatment with any of the compounds tested (supplemental table 4). The inability of these drugs to address the root cause of the disease, along with the poor translatability of the *fmr1* KO mouse model, could have contributed to their lack of efficacy. Future studies should also determine if this threshold level of FMRP is consistent across multiple patient lines with different CGG repeat expansions and methylation statuses, as well as to elucidate the levels of FMRP required to correct other functional deficits such as synaptic scaling.

In summary, we have demonstrated an FMRP-dependent neuronal phenotype of increased spontaneous activity in FXS iPSC-derived neurons that can be normalized through partial restoration of FMRP. Our data suggest that either greater than 20% FMRP-expressing neurons in a mosaic population, or greater than 5% overall FMRP expression levels throughout a neuronal network can be used as thresholds for evaluating potential therapeutic interventions focused on reactivating FMRP in iPSC-derived neurons from FXS patients.

## Author Contributions

J.G., H.W., and A.C. conceived the idea for this project. J.G. and H.W. designed the experiments and interpreted the data. J.G., H.W., C.N., C.S., V.V., D.Q. and K.J. performed the experiments. R.J. provided dCas9-Tet1, dCas9-dTet1 and FMR1 CGG gRNA constructs. S.W. provided the FXS_SW isogenic iPSCs. J.G. and H.W. wrote the manuscript with inputs from all the other authors.

## ACKNOWLEDGMENTS

We thank Drs. Shawn Liu and Rudolf Jaenisch at Whitehead Institute at MIT for dCas9-Tet1, FRM1 CGG plasmids, and pX458 Cas9-T2A-GFP gRNA backbone plasmid, Dr. Steve Warren at Emory University for providing FXS_SW isogenic iPSC lines, Dr. Marius Wernig at Stanford University for providing pTet-O-NGN2-2A-Puro plasmid. We thank members of Fulcrum Therapeutics for discussion and suggestions on the manuscript. R.J. is co-founder of Fate Therapeutics, Fulcrum Therapeutics and Omega Therapeutics.

---

**Supplemental Table 1.**

| **iPSC cell lines** | **Reference** |
| --- | --- |
| FXS_SW | Xie et al., 2016 |
| C1_2_SW | Xie et al., 2016 |
| Epi49 WT | ThermoFisher Scientific, A18945 |
| FMR1 KO | this study |

**Supplemental Table 2.**

| **Taqman probe name** | **Vendor** | **Assay ID** | **Cat. No.** |
| --- | --- | --- | --- |
| FAM-FMR1 | ThermoFisher Scientific | Hs00924547_m1 | 4331182 |
| POP4-ABY | ThermoFisher Scientific | Hs00198357_m1qsy | CCU002NR |

**Supplemental Table 3.**

| **<u>Antibodies</u>** | **<u>Vendor</u>** | **<u>Cat. No.</u>** | **<u>Dilution</u>** |
| --- | --- | --- | --- |
| Mouse anti FMRP (5C2) | BioLegend | 834701 | 1:200 |
| Mouse anti Cas9 (7A9-3A3) | Active Motif | 61577 | 1:100 |
| Chicken anti TUJ1 | AVES | TUJ89947984 | 1:500 |
| DAPI | Sigma-Aldrich | 10236276001 | 1:2000 |
| Donkey anti-Mouse IgG (H+L) | Invitrogen | A-21203 | 1:2000 |
| Goat anti-Chicken IgY (H+L) | Invitrogen | A-11039 | 1:2000 |
| Donkey anti-Mouse IgG (H+L) | Invitrogen | A-31571 | 1:2000 |

**Supplemental Table 4.**

| **FMR1 Relative Expression** |  |  |  |
| --- | --- | --- | --- |
| **<u>Treatment</u>** | **<u>Avg (% Control)</u>** | **<u>StDev</u>** | **<u>n</u>** |
| *Control (DMSO)* | 100.00 | 20.51 | 3 |
| *FXS (DMSO)* | 0.08 | 0.02 | 3 |
| *Baclofen* | 0.44 | 0.09 | 3 |
| *5-AzaC* | 2.75 | 0.74 | 3 |
| *Rimonabant* | 0.26 | 0.03 | 3 |
| *Lovastatin* | 2.58 | 4.54 | 3 |
| *Metformin* | 0.55 | 0.13 | 3 |
| *Mavoglurant* | 0.60 | 0.09 | 3 |
| *MPEP* | 0.22 | 0.01 | 3 |
| *Valproic Acid* | 0.46 | 0.10 | 3 |
| *Minocycline* | 0.16 | 0.01 | 3 |
| *Splitomicin* | 0.08 | 0.07 | 3 |

